# Overexpression of IκBα modulates NF-κB activation of inflammatory target gene expression

**DOI:** 10.1101/2023.03.14.532132

**Authors:** Polly Downton, James S Bagnall, Hazel England, David G Spiller, Neil Humphreys, Dean A Jackson, Pawel Paszek, Michael R H White, Antony D Adamson

## Abstract

Cells respond to inflammatory stimuli such as cytokines by activation of the nuclear factor-κB (NF-κB) signalling pathway, resulting in oscillatory translocation of the transcription factor p65 between nucleus and cytoplasm to mediate immune response. We investigate the relationship between p65 and inhibitor-κBα (IκBα) protein levels and dynamic properties of the system, and how this interaction impacts on the expression of key inflammatory genes. Using bacterial artificial chromosomes, we developed new cell models of IκBα-eGFP protein overexpression in a native genomic context. We find that cells with high levels of the negative regulator IκBα remain responsive to inflammatory stimuli and maintain dynamics for both p65 and IκBα. In contrast, canonical target gene expression is dramatically reduced by overexpression of IκBα, but can be partially rescued by overexpression of p65. Treatment with leptomycin B to promote nuclear accumulation of IκBα also suppresses canonical target gene expression, suggesting a mechanism in which nuclear IκBα accumulation prevents productive p65 interaction with promoter binding sites. This causes reduced target promoter binding and gene transcription, which we validate by chromatin immune precipitation and in primary cells. Overall, we show how inflammatory gene transcription is modulated by the expression levels of both IκBα and p65, and that transcription can be partially decoupled from p65 protein dynamics. This results in an anti-inflammatory effect on transcription, demonstrating a broad mechanism to modulate the strength of inflammatory response.

## Introduction

The nuclear factor kappa B (NF-κB) signaling pathway is involved in the regulation of a wide range of cellular processes. NF-κB signalling is a key mediator of the immune system, and is activated in many cell types in response to viral and bacterial pathogens and cytokine signalling cascades (Hayden and Ghosh, 2008). Dysregulated NF-κB signalling is linked to cancer, inflammatory and autoimmune diseases (Perkins, 2012, Taniguchi and Karin, 2018, Barnabei et al., 2021).

In mammals there are five NF-κB proteins; p65/RelA, RelB, cRel, p50 and p52, which hetero- and homo-dimerise in several possible combinations (Hayden and Ghosh, 2008). NF-κB transcription factor complexes (canonically, RelA/p65:p50) are found in the cytoplasm of resting cells bound to inhibitor κB (IκB) family molecules, the most abundant of which is IκBα. Stimulation of cells with a pro-inflammatory signal, such as tumour necrosis factor α (TNFα), leads to a signalling cascade which results in IκB kinase (IKK) activation, and phosphorylation of both p65 and IκBα. The phosphorylated IκBα is subsequently ubiquitinated and targeted for proteasomal degradation, releasing p65 to translocate to the nucleus and bind to target gene promoters, including IκBα. Newly synthesised IκBα can translocate to the nucleus, where it may bind p65 to result in relocation of the p65:IκBα complex to the cytoplasm.

We and others have shown that p65 oscillates between nucleus and cytoplasm in the presence of continued stimulation. This oscillatory behaviour quickly becomes asynchronous in a population of cells following initial stimulation, meaning single cell analyses are essential for detection and quantification (Paszek et al., 2010, Aqdas and Sung, 2022). Exogenous plasmid and lentiviral systems typically include a constitutive promoter to drive coding sequence expression, which can result in perturbation of transcript copy number and protein expression level. It has been shown that overexpression of p65 results in downstream effects on target gene expression (Sung et al., 2009, Lee et al., 2014), often resulting in increased activation of pro-inflammatory target genes.

We previously generated a clonal cell line containing a stably-integrated recombinant Bacterial Artificial Chromosome (BAC) expressing IκBα fused to eGFP, under the regulatory control of its native genomic context (Adamson et al., 2016). We showed that the oscillatory timing of IκBα expression is robust and out-of-phase with p65 nuclear:cytoplasmic (N:C) translocation. We identified a heterogeneous refractory period of response to pulsatile cytokine treatment, predicted to be controlled by a post-translational switch between the ligand activated receptor and the IKK, which determines whether cells can respond to a second pulse of cytokine stimulation.

The delivery of the IκBα-eGFP BAC to cells and successful stable integration into genomic DNA resulted in overexpression of IκBα. Previous observations have shown that p65 protein level has an effect on NF-κB target gene expression (Sung et al., 2009). We investigate the effect of IκBα feedback on oscillatory behaviour and downstream gene expression using an overexpression system. We find, using BAC stable cell lines and cells derived from transgenic mice, that although oscillatory period is robust to differing expression levels, IκBα overexpression results in potent downregulation of the transcription of canonical NF-κB target genes. We provide evidence of a mechanism relating this to increased levels of free IκBα in the nucleus, where IκBα directly competes for translocated p65.

## Materials & Methods

### Reagents and cell culture

SK-N-AS neuroblastoma (Cat. No. 94092302) cells were obtained from the European Collection of Authenticated Cell Cultures (ECACC). Cells were cultured according to ECACC protocols, and frozen down to form a low passage working stock. Working stocks were screened to ensure the absence of mycoplasma every 3 months using LookOut Mycoplasma PCR Detection Kit (Cat. No. D9307 Sigma, UK). Cells were cultured in Modified Eagle’s Medium supplemented with 10% foetal bovine serum (FBS) and 1% non-essential amino acids. Treatments used recombinant human TNFα (10 ng/ml; Merck 654245) and/or LMB (20 ng/ml, Merck 431050).

Primary mouse fibroblasts were cultured from ear tissue biopsies. Tissue was minced and cells isolated by incubation in collagenase (Sigma C2674) for 30 minutes at 37°C. Cells were maintained in Dulbecco’s Modified Eagle’s Medium supplemented with 10% FBS and penicillin/streptomycin. Cells were treated with recombinant mouse TNFα (10 ng/ml, Merck 654245).

### IκBα-eGFP construct

We have previously described the IκBα-eGFP recombinant BAC (Adamson et al., 2016). Briefly, using a GalK selection/counterselection recombineering strategy (Warming et al., 2005) we seamlessly integrated the fluorescent protein genes in place of the STOP codons, to create fusion constructs that are expressed in a pseudo-genomic context when transfected/integrated into cells. This construct is available on request.

### Generation of IκBα-eGFP clonal cell lines

BAC DNA for transfection was prepared using the BAC100 Nucleobond kit (Macherey-Nagel, Germany). Cells were transfected using ExGen500 transfection reagent (Fermentas, UK) and clonal cell lines were derived by cell sorting as previously described (Adamson et al., 2016). Cell lines are available on request.

### Lentiviral packaging and transduction

Lentiviral constructs were cloned and lentivirus produced as previously described (Bagnall et al., 2015). Clonal cell lines were transduced with lentivirus encoding human p65-mCherry under control of the ubiquitin ligase C promoter.

### qRT-PCR

For RNA isolation, cells were seeded into 6-well plates (100000/well). After treatments as described, cells were washed once with cold PBS then lysed. Total RNA was isolated using the High Pure RNA isolation kit (Roche). RNA concentration was determined using a Nanodrop ND-1000 spectrophotometer (Thermo). RNA was reverse transcribed to cDNA using the SuperScript VILO cDNA synthesis kit (Life Technologies). The resulting cDNA was analysed by qRT-PCR on a LightCycler 480 using SYBR Green 1 Master Mix (Roche). Relative fold change in expression was determined by the ddCt method, using *PPIA* expression as a housekeeping control. Primer sequences are given in Table 1.

### smRNA-FISH

Custom smRNA-FISH probe sets were designed against coding sequences, and UTRs when necessary, using the Stellaris FISH Probe Designer (Biosearch Technologies Inc). Probes were conjugated with Quasar 570 or Quasar 670. Sequences are given in Table 2. *eGFP* transcripts were detected using the pre-designed Quasar 570 probe set (Biosearch Technologies Inc., VSMF-1014-5).

Cells were seeded into 12-well plates (40000/well) containing coverslips pre-coated with poly-L-lysine. Following treatment, cells were washed, fixed with 3.7% formaldehyde in PBS for 10 minutes, then permeabilised with 70% ethanol for 2-24 hours at 4°C. Coverslips were washed (10% formamide in 2 X SSC) then hybridised with probe mix (probe/s of interest in 10% formamide in 2 X SSC containing 100 mg/ml dextran sulphate) overnight at 37°C. Coverslips were washed, incubated with DAPI, then mounted in Vectashield for imaging.

Images were acquired on a DeltaVision (Applied Precision) microscope using a 60x NA 1.42 Plan Apo N objective and a Sedat Quad filter set. The images were collected using a CoolSNAP HQ (Photometrics) camera with z optical spacing of 0.2 μm. Raw images were deconvolved using softWoRx software. Deconvolved image stacks were analysed using FISH-quant to determine transcript numbers per cell (Mueller et al., 2013). Exemplar maximum projection images were generated in Fiji (Schindelin et al., 2012).

### Western blotting

Cells were seeded in 35 mm dishes (50000/dish) two days prior to sample collection. Samples were washed once with cold PBS, then lysed in hot buffer (1% (w/v) SDS, 10% (v/v) glycerol, 10% (v/v) b-ME, 40 mM Tris pH 6.8, 0.01% (w/v) bromophenol blue). Proteins were resolved on polyacrylamide gels run under denaturing conditions, then transferred to nitrocellulose membrane (Protran BA-83, GE Healthcare). Membranes were blocked with 5% (w/v) skim milk powder in TBS-T prior to overnight incubation with primary antibody (anti-IκBα, Cell Signalling Technology #9242, 1:1000; anti-ICAM1, Santa Cruz Biotechnology sc-8439, 1:200; anti-α-Tubulin, Sigma Aldrich T6199, 1:4000; anti-Vinculin, Cell Signalling Technology 4650S, 1:1000). Membranes were washed with TBS-T, incubated with HRP-conjugated secondary antibody (CST #7074 or #7076, 1:1000), washed and developed using Luminata Crescendo substrate (Millipore WBLUR0500). Signal was detected using Carestream Kodak BioMax MR film (Sigma-Aldrich). Unprocessed images of films are provided in Figure S5. Signal was quantified using Fiji (Schindelin et al., 2012).

### Confocal time lapse imaging

Cells were seeded into 35 mm glass-bottomed dishes (Greiner) and imaged using several Zeiss confocal microscopes (LSM Pascal, Exciter, 710, 780, 880) with Fluar 40x NA 1.3 objectives. Cells were maintained at 37°C in humidified 5% CO_2_ throughout image acquisition. Image capture used Zeiss software (Aim version 4.2, Zen 2010b SP1 or Zen 2.1 SP3 FP2). Quantification of IκBα-eGFP fluorescent signal of whole cells was performed using region of interest (ROI) analysis in Zen 2010b SP1 software. Normalised expression level was calculated relative to average cell fluorescence intensity prior to treatment. Quantification of dynamic parameters was performed by a custom script based on the ‘findpeaks’ function using Matlab 2020a. Exemplar image sequences were generated in Fiji (Schindelin et al., 2012).

### Fluorescence Correlation Spectroscopy (FCS)

Cells were seeded into 35 mm glass-bottomed dishes and imaged using a Zeiss LSM 880 microscope, as described above Fluorescence fluctuations were recorded in five separate measurements of 5 seconds for manually selected discrete locations in either the cytoplasm or nucleus across many cells, with the pinhole set to measure 1 AU, the equivalent of 0.75 fL volume when using 488 nm laser. Data was analysed using the ‘Fish-and-Cushion’ software as described in (Koch et al., 2022). This performs autocorrelation analysis, which determines the concentrations of fluorescent protein by selecting parameters from the best fit model across a range of models that captures protein mobility and photochemistry fluctuations for each cell measurement.

### Nanostring

RNA was isolated as described above. Samples were analysed with the Nanostring nCounter analysis system and a custom CodeSet (Table 3). Data was processed in nSolver Analysis Software v4.0, with normalisation to five housekeeping genes and internal positive control probes. Clustering analysis grouped genes based on Pearson correlation coefficient of log count values, with linkage calculated from average distance between elements.

### ChIP

Cells were seeded into 150 mm dishes (3M/dish) and allowed to grow until near confluent. Cells were treated with TNFα+/- LMB as described, then processed using the EZ-Magna ChIP kit (Millipore 17-409). Chromatin was fragmented by sonication at 4°C using a Bioruptor 300 (Diagenode; 45 cycles, 30s on/30s off). Immunoprecipitation used 4 μg ChIP validated antibody against NF-κB p65 (Millipore, 17-10060), or control antibodies against polII or IgG. Chromatin immunoprecipitation was quantified by qPCR.

### Animals

Mice were maintained in the University of Manchester Biological Services Facility. All protocols were approved by the University of Manchester Animal Welfare and Ethical Review Body and licenced under the Animals (Scientific Procedures) Act 1986.

### Generation of transgenic mice

To create BAC transgenic mice 10 μg maxi-prepped BAC DNA (Nucleobond 100) was linearised by restriction digest (NotI) and purified by sepharose column purification (GE Healthcare). Briefly, a standard 5 ml pipette was used as a column, with the cotton plug removed and filled with injection buffer (sterile filtered 10 mM Tris (pH 7.5), 0.1 mM EDTA (pH 8.0), 100 mM NaCl) equilibrated sepharose beads. Linearised DNA, with bromophenol blue dye, was added to the column, and once the dye had entered the beads more injection buffer added. Fractions were collected every five minutes until dye had drained from the column. A sample of each fraction was run on an agarose gel to confirm DNA purity, and the DNA containing fraction was diluted to 2 ng/μl for mouse zygote injection.

Zygote injections, in C57/BL6j background embryos, were performed by the Manchester Genome Editing Unit. Pups were genotyped using primers specific to the fluorescent protein gene. The strain is cryopreserved and available on request.

### Targeted Locus Amplification genotyping

Mice aged 6-8 weeks were culled by an S1 method. The spleen was removed and prepared as described in the Cergentis spleen sample preparation protocol. Samples were sent to Cergentis (Utrecht, Netherlands) for identification of transgene integration site and copy number analysis.

### Statistical analysis

Statistical analysis used GraphPad Prism. Details of sample size and statistical tests are provided in figure legends.

## Results

### Generation and characterisation of clonal SK-N-AS IκBα-eGFP BAC cell lines

We have previously described the generation of a recombinant BAC expressing an IκBα-eGFP fusion protein in its native genomic context (Adamson et al., 2016). This includes over 150 kb of flanking, intronic and UTR sequence to ensure gene expression is subject to natural regulatory processes and feedbacks. We generated a series of clonal SK-N-AS neuroblastoma cell lines by single cell sorting and antibiotic selection (Figure 1A), and identified two clones, here termed IκBα A and IκBα B (previously named clone C9, (Adamson et al., 2016), with elevated IκBα expression levels. In order to visualise p65 in these backgrounds, these clones were also transduced with a lentiviral p65-mCherry expression vector (Figure 1A).

**Figure 1.**
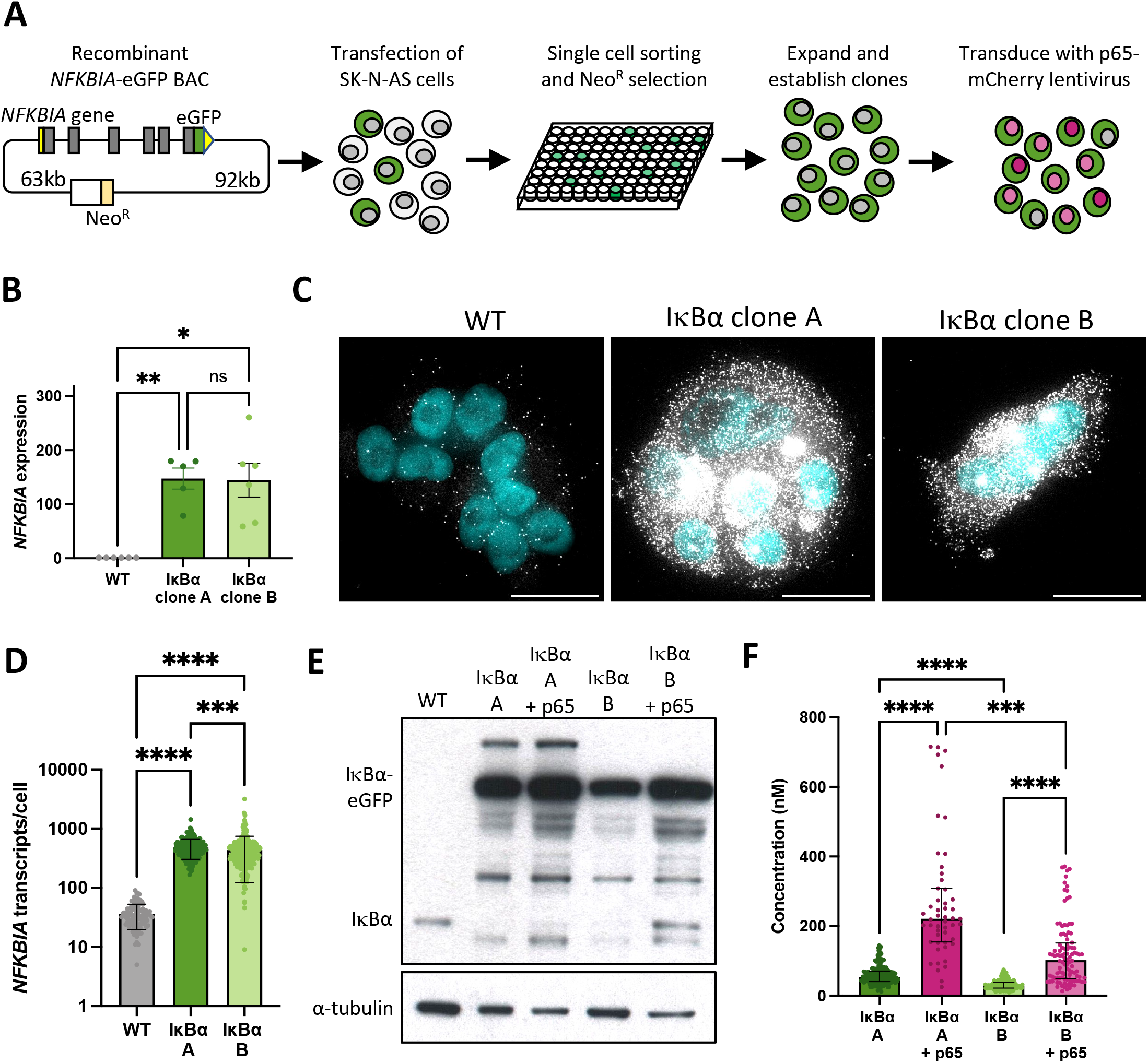
Generation and characterisation of IκBα-eGFP reporter cells. A: Schematic representation of cell line generation. Clonal cell lines with integration of a BAC expressing an IκBα-eGFP fusion protein were selected. Validated clonal cell lines were subsequently transduced with p65-mCherry lentivirus. B: qRT-PCR measurement of *NFKBIA* gene expression in unstimulated conditions. Expression is normalised to *PPIA*. N = 5-6, One-way ANOVA, Kruskal-Wallis test, Dunn’s multiple comparison correction. C: smRNA-FISH detection of *NFKBIA* transcripts in unstimulated conditions. Probe is shown in white; nuclei counterstained with DAPI are shown in cyan. Scale bar = 20 µm. D: Quantification of *NFKBIA* by smRNA-FISH detection in clonal cell lines in unstimulated conditions. N = 100-200 cells, imaged over 2-3 independent experiments. One-way ANOVA, Kruskal-Wallis test, Dunn’s multiple comparison correction. E: Protein quantification by Western blot in clonal cell lines in unstimulated conditions. F: Cytoplasmic protein quantification by FCS in clonal cell lines in unstimulated conditions. N = 50-200 cells, imaged over 3-4 independent experiments. Error bars indicate median +/- interquartile range. One-way ANOVA, Kruskal-Wallis test, Dunn’s multiple comparison correction. Throughout, error bars indicate mean +/- SD unless otherwise stated; * indicates p < 0.05, ** indicates p < 0.01, *** indicates p < 0.001, *** indicates p < 0.0001.

We used a range of approaches to quantify IκBα expression in these clones. Quantitative RT-PCR indicates levels of IκBα transcript far in excess of WT cells (Figure 1B). Using probes directed against IκBα we identified between 10-100 IκBα transcripts per cell in unmodified (WT) SK-N-AS cells by single molecule RNA fluorescence *in situ* hybridisation (smFISH; Figure 1C). In comparison, we detected between 100-1000 transcripts per cell in the BAC clones (mean of 480 transcripts per cell in clone A, 434 in clone B; Figure 1D), typically densely clustered around a single bright spot, representing the site of active transcription from the integrated BAC constructs. This may be an underestimation of transcript number since saturated signal detection at the sites of transcription meant transcripts could not be individually resolved at these sites. Western blot analysis of protein extracts revealed elevated IκBα-eGFP levels in both clones, with and without p65-mCherry (Figure 1E). We detect a number of additional protein bands which may represent degraded or intermediate products in the overexpression cell lines. We also observed a reduction in the level of endogenous, untagged IκBα protein in both overexpression clones. Finally, we used Fluorescence Correlation Spectroscopy (FCS) to measure the intracellular concentration of IκBα-eGFP. In IκBα clone A, FCS determined the mean cytoplasmic concentration of IκBα-eGFP to be 59 nM (+/- 26 nM), compared to 32 nM (+/- 13 nM) in IκBα clone B. In both clones, co-overexpression of p65-mCherry resulted in increased expression of IκBα-eGFP (Figure 1F), corroborating previous studies (Lee et al., 2014). These approaches all confirm that clones A and B exhibit elevated IκBα levels compared to WT cells, with Clone A showing higher protein expression level than Clone B.

### IκBα oscillation dynamics are independent from IκBα expression level

IκBα is an early target gene responsive to stimulation by TNFα treatment (Hao and Baltimore, 2009). We found overall levels of IκBα transcript in our BAC clonal cell populations to be elevated even in the absence of TNFα stimulation (Figure 1), so next we used smFISH to characterise the effect of TNFα stimulation on transcript levels in single cells (Figure S1A). In untreated WT cells we detected a low background level of cytoplasmic IκBα mRNA, which increased significantly after 130’ TNFα treatment (Figure S1A, upper panels, and Figure S1B). One or two distinct bright signal spots, indicating the allelic sites of transcription, are seen in the nuclei of treated cells (Figure S1A, upper panels). Treatment of cell lines with TNFα resulted in detection of extremely high transcript levels, suggesting further induction of expression over the high basal level in untreated clonal cells (and prohibiting accurate quantitative analysis of transcript numbers). Co-staining for *eGFP* transcript sequences confirms that most cellular transcripts are derived from the integrated BAC *NFKBIA-eGFP* transgene. This indicates the integration of IκBα-eGFP BACs results in significant overexpression of IκBα transcript when compared to WT cells, in both untreated and treated conditions.

Our previous studies have found p65 oscillates from cytoplasm to nucleus in the continuous presence of pro-inflammatory stimuli such as TNFα (Nelson et al., 2004, Ashall et al., 2009). We have also shown, through mathematical modelling and single cell imaging of cells transfected with the IκBα-eGFP BAC, that IκBα degrades and resynthesizes out-of-phase to p65 nuclear movements, as expected in a negative feedback regulatory loop (Nelson et al., 2004, Adamson et al., 2016). IκBα oscillations have been observed in several cell types tested, including primary cells derived from transgenic mice (Harper et al., 2018). We next examined the dynamic inflammatory response of our stable cell lines to TNFα by live cell confocal microscopy. At the protein level, continuous TNFα treatment resulted in rapid degradation of IκBα-eGFP in both clones (median trough around 30 min), followed by resynthesis of IκBα-eGFP with an initial peak 90-110 minutes after treatment (Figures 2A and 2B). Cells continued to show oscillations in fluorescence intensity as a result of IκBα-eGFP degradation and synthesis, quickly becoming asynchronous (Supplementary Video 1). Response to a short pulse of TNFα demonstrates that IκBα degradation and resynthesis rates are consistent between cell lines, and comparable to unmodified cells (Figure S2). Analysis of oscillatory behaviour in clones transduced with p65-mCherry confirmed p65 nuclear translocation occurs out-of-phase with IκBα degradation/resynthesis cycles (Figure 2C), as seen previously (Adamson et al., 2016). Median oscillatory period in both BAC cell lines was circa 110 minutes (Figure 2D). This corroborates data previously generated using other NF-κB expression systems, including overexpression of p65 fusions from constitutively active promoters in exogenous vectors (Adamson et al., 2016, Son et al., 2022, DeFelice et al., 2019), from endogenously tagged alleles in MEFs derived from an GFP-p65 transgenic mouse (Zambrano et al., 2016), and from a CRISPR targeted allele in MCF7 cells (Stewart-Ornstein and Lahav, 2016). Analysis of clones transduced with p65-mCherry found an increased amplitude of oscillation, with broader peak width and slightly increased median average period of 130 minutes (Figure 2D-F). This is consistent with previous work, which found that cells maintain a robust oscillatory period of 100-110 minutes when IκBα or p65 are overexpressed in isolation, but that overexpression of p65 and feedback-responsive IκBα results in lengthening of the oscillatory period (Nelson et al., 2004). Previous experiments with IκBα under a non-native 5xNF-κB response element promoter resulted in dramatic lengthening of the period to around 200 minutes; however, this system is unlikely to reflect all regulatory inputs as effectively as the pseudo-native context provided by our BAC reporter system.

**Figure 2.**
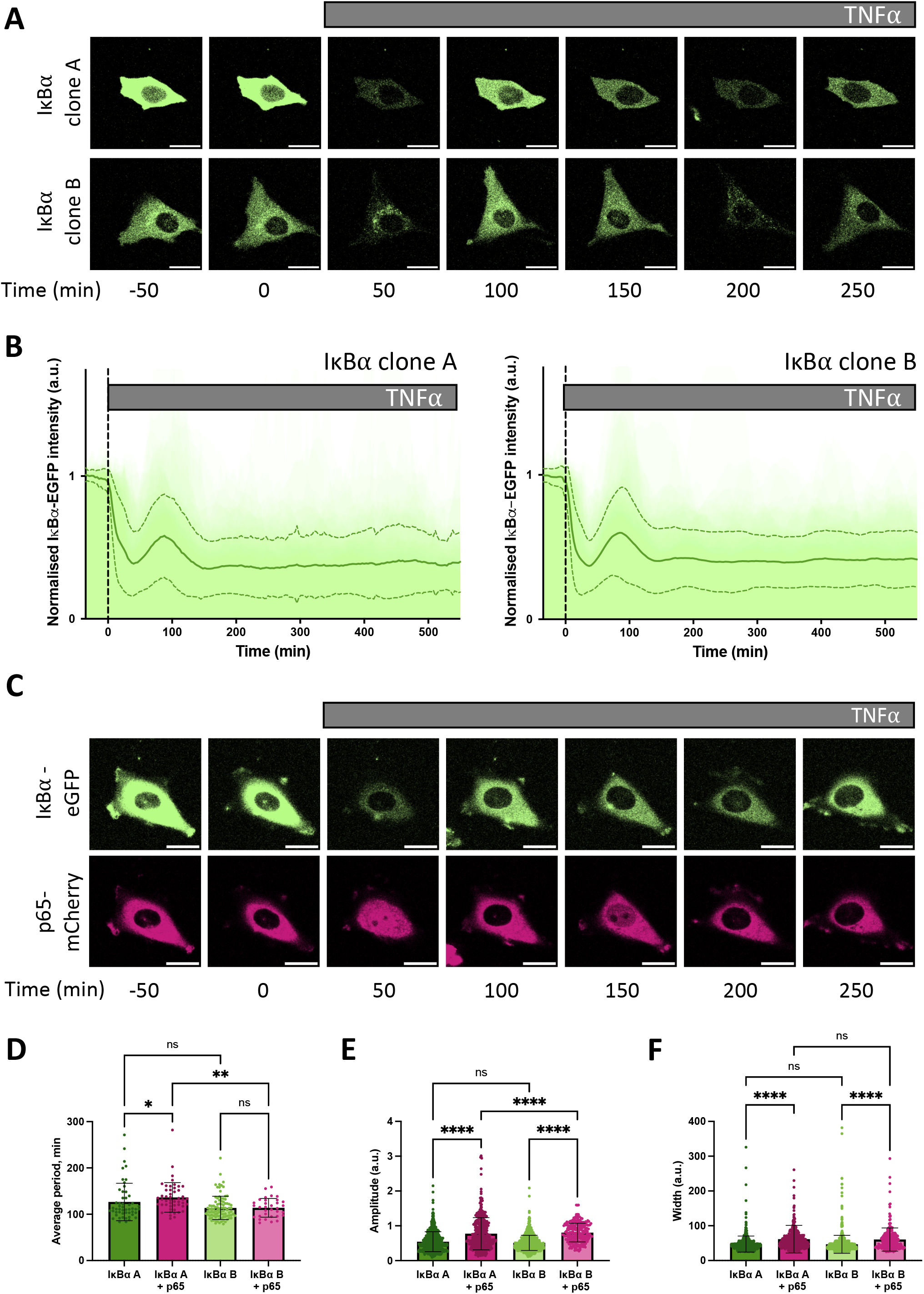
Dynamic response of IκBα-eGFP reporter cells to TNFα treatment. A: Example time course images of IκBα-eGFP clonal cell response to TNFα treatment. Scale bar = 20 µm. B: Clonal cell response to continuous TNFα treatment. Individual traces are shown in pale green; population average +/- SD is shown in dark green. n = 140-160 cells imaged over at least 6 independent experiments. C: Example time course images of IκBα-eGFP clonal cells transduced with p65-mCherry lentivirus response to TNFα treatment. Scale bar = 20 µm. D, E, F: Quantification of dynamic cell response to continuous TNFα treatment. Normalised fluorescence intensity was tracked over at least three oscillation cycles and traces quantified to determine average period of oscillation (D), peak amplitude (E) and peak width (F). N = 30-90 cells/clone imaged over at least 2 independent experiments. One-way ANOVA, Kruskal-Wallis test, Dunn’s multiple comparison correction.

### IκBα overexpression modulates NF-кB target gene transcription

We and others have shown an association between oscillatory behaviour of the NF-κB signalling system and downstream gene expression (Ashall et al., 2009, Martin et al., 2020). Given overexpression of IκBα did not perturb oscillatory behaviour, we investigated whether expression of NF-кB target genes was affected. Wild type (WT) SK-N-AS cells, IκBα Clone A, IκBα Clone B and IκBα Clone B + p65-mCherry transduced cells were treated with TNFα and RNA extracted over a time course. Transcript abundance for a pre-defined NF-кB target gene subset was quantified by Nanostring analysis (Supplementary Table 1). Hierarchical cluster analysis broadly grouped target gene response into four patterns of behaviour (Figure S3). The largest cluster, Cluster 1, includes prototypical inflammatory mediators and NF-κB feedback genes which show sustained activation over 430 minutes in unmodified SK-N-AS cells (Figure 3A). In IκBα clone A cells, TNFα treatment resulted in substantially reduced activation of these gene targets. IκBα clone B cells showed a strong dampening of activation, which could be partially rescued by co-overexpression of p65-mCherry in lentivirus transduced cells. Genes in Cluster 3 showed basal upregulation in response to IκBα overexpression in the clonal cell lines, whilst genes in other clusters demonstrated inter-clone variability. We focused on genes from Cluster 1 for further investigation.

**Figure 3.**
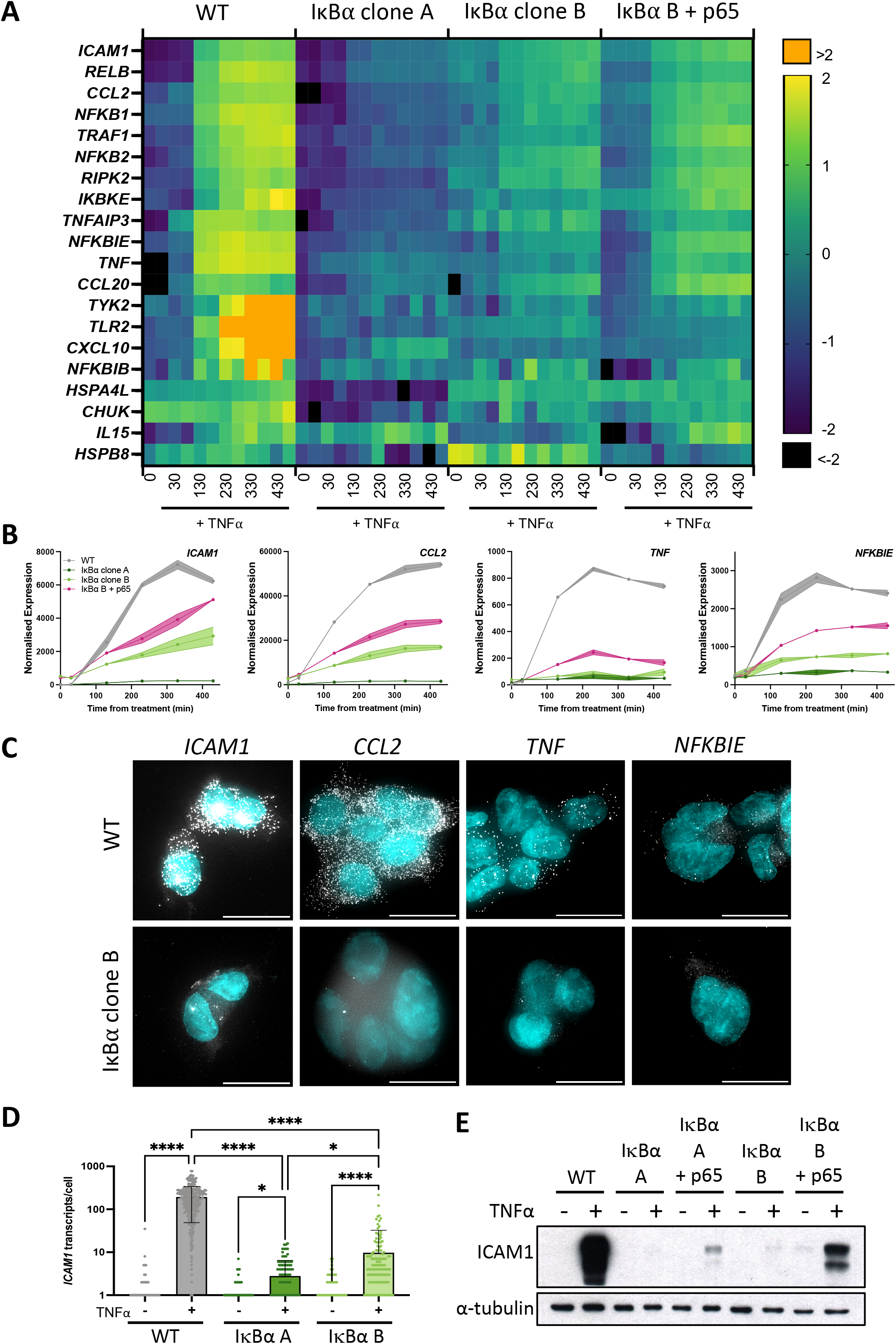
Overexpression of IκBα suppresses target gene expression. A: Heatmap representation of gene expression of clonal cell lines in response to TNFα treatment, as determined by Nanostring gene expression assay. N = 2/time/cell line. Expression is normalised to five housekeeping genes and internal control probes, then scaled to average gene expression level. B: Normalised Nanostring counts for selected genes from (A), average +/- SD. C: Example images showing smRNA-FISH detection of target gene transcripts in cell lines after 130 min TNFα treatment. Scale bar = 20 µm. D: Quantification of *ICAM1* transcript by smRNA-FISH in cell lines before and after treatment with TNFα for 130 min. N = 90-300 cells/condition, imaged over six independent experiments. One-way ANOVA, Kruskal-Wallis test, Dunn’s multiple comparison correction. E: Quantification of ICAM1 protein expression before and after treatment with TNFα treatment for 230 min.

We validated the Nanostring results using complementary approaches for several genes showing behaviour typical of Cluster 1: *ICAM1, CCL2, TNF* and *NFKBIE* (Figure 3B). These genes have been characterised as direct targets of NF-κB signalling and are upregulated in response to inflammatory cytokine treatment (Astarci et al., 2012, Sutcliffe et al., 2009, Ghosh et al., 2010). Transcript level of target genes was quantified by smFISH, following treatment with TNFα for 130 minutes (Figure 3C; treated cells for WT and IκBα clone B shown). Consistent with the gene expression data from Nanostring analysis, WT cells exhibited an increase in transcript number after treatment, but IκBα clone B cells showed comparatively low transcript levels for all gene targets. In common with previous studies, numbers of transcripts per cell showed considerable heterogeneity for some targets e.g. *TNF* (Bass et al., 2021, Bagnall et al., 2018, Bagnall et al., 2020). Quantification of transcript number found WT cells express 176-210 *ICAM1* transcripts per cell after 130 minutes TNFα stimulation (95% confidence interval, CI), whilst IκBα clone A expressed an average of 3 (95% CI 2-4) transcripts per cell, and clone B expressed an average of 10 (95% CI 6-13) transcripts per cell (Figure 3D). Western blot analysis confirmed strong protein expression of ICAM1 in TNFα-treated WT cells, but undetectable and near-undetectable levels of ICAM1 protein in clones A and B respectively (Figure 3E). Again, the expression response could be partially rescued by co-overexpression of p65-mCherry (Figure 3E). These data indicate that whilst IκBα protein oscillations and p65 N:C translocations are robust to changes in IκBα level, overexpression of IκBα can result in profound effects on target gene activation. NFκB activity is regulated by multiple feedback loops, and previous work has hypothesised IκBα feedback dictates the amplitude of response, with other feedbacks (e.g. A20) altering timing (Fagerlund et al., 2015, Adamson et al., 2016). We find that, In our system, response timing is unaltered but the ‘amplitude’ of response (i.e. activation of target genes) has been dampened by IκBα overexpression.

### IκBα-eGFP overexpression leads to elevated nuclear IκBα and competition with target DNA response elements for nuclear p65

The correlation between elevated IκBα expression and canonical target gene repression, together with partial gene expression rescue by increased p65 concentration, indicates these effects are likely to be a response to variation in intracellular protein concentration. Mechanistic modelling has previously found that sensitivity to TNFα is dependent upon the cellular ratio of NFκB to its inhibitor, and nucleocytoplasmic shuttling of these complexes in resting cells (Patel et al., 2021). IκBα has both nuclear import and export sequences, and the equilibrium between these two processes determines its net distribution.

We and others have previously shown that IκBα accumulates in the nucleus upon cellular treatment with the nuclear export inhibitor Leptomycin B (LMB) (Nelson et al., 2004, Rodriguez et al., 1999) (Sung et al., 2009), indicating that subcellular localisation of IκBα is dynamic. Such an equilibrium has previously been found for NFκB: IκBα complexes (Huang et al., 2000); we hypothesise that under basal conditions a similar equilibrium exists between IκBα nuclear export and import in our cells. This results in an elevated basal concentration of nuclear IκBα in our overexpressing cells. To confirm this, we used FCS to measure the concentration of fluorescent IκBα-eGFP in the nucleus of the BAC clonal cell lines (Figure 4A). We detected a nuclear presence of IκBα in both clones, with a slightly higher concentration for IκBα clone A (25 nM vs 16 nM). Treating cells with LMB to block nuclear export resulted in accumulation of IκBα-eGFP in the nucleus of both clones (Figure 4B). The rate of import was similar in each clone (Figure 4C), indicating this rate is unaffected by IκBα expression level. These data suggest the proportion of IκBα molecules in the cytoplasm vs. nucleus of a given cell is correlated (as previously observed in (Kardynska et al., 2018)), and nuclear import/export maintains this equilibrium, resulting in nuclear concentration of IκBα being a fixed proportion of total cellular IκBα. As clone A cells have higher overall IκBα protein expression levels (Figure 1), this results in higher nuclear IκBα levels in comparison to clone B (Figure 4A). Thus, in our system, the BAC-mediated increase in IκBα expression results in an increase in nuclear IκBα concentration.

**Figure 4.**
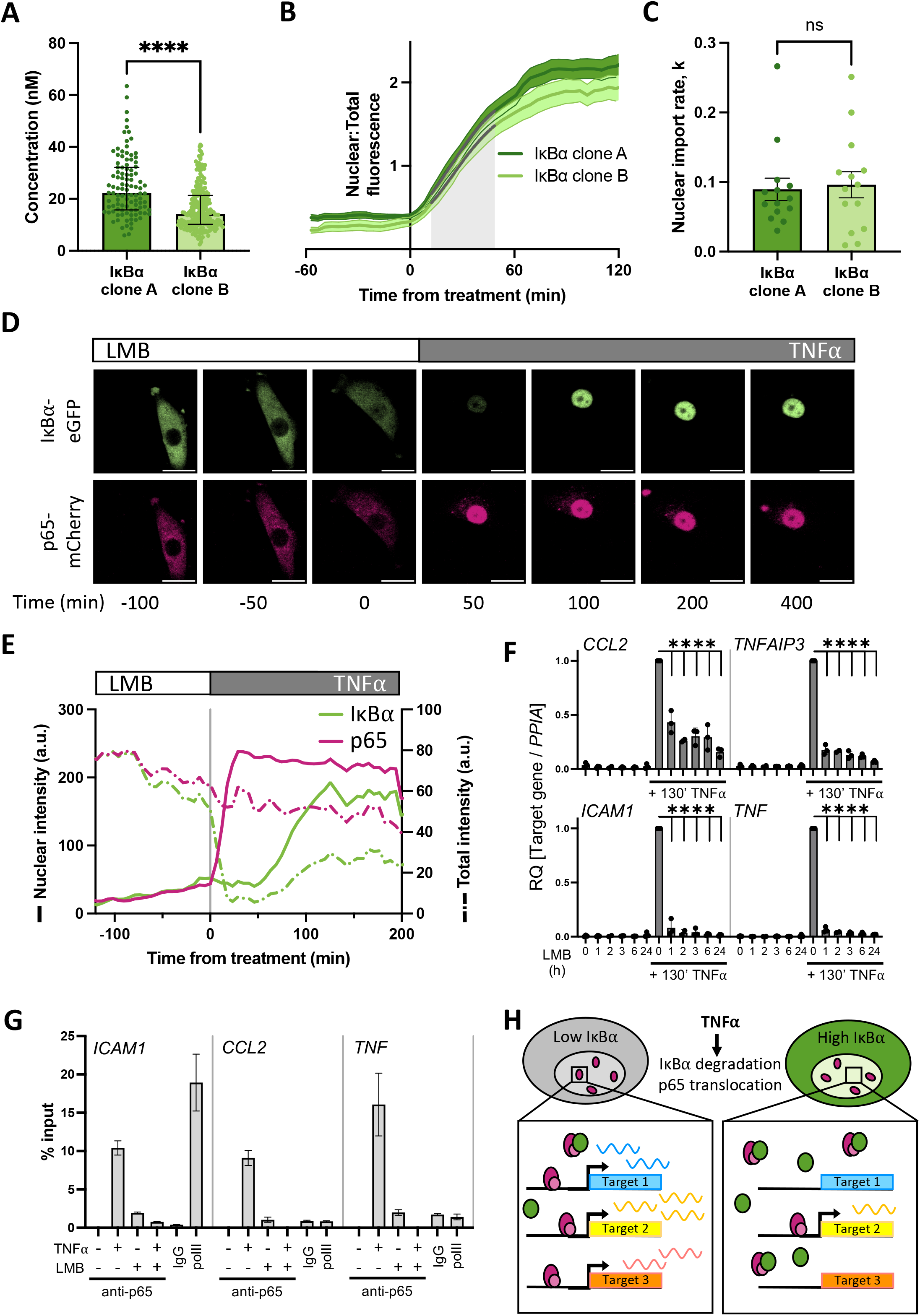
Nuclear accumulation of IκBα suppresses target gene expression. A: FCS quantification of nuclear protein in clonal cell lines in unstimulated conditions. Error bars indicate median +/- interquartile range. Mann-Whitney test, N = 100-250 cells over 3 independent experimental replicates. B: Quantification of nuclear fluorescence (average +/- SD) following LMB treatment of cells. Import rate was calculated from 10 to 50 minutes after treatment (shaded grey) from N = 14 cells/clone. C: IκBα nuclear import rate following LMB treatment, calculated from nuclear fluorescence data in (B) using a non-linear model (model fit indicated by grey lines on B). Mann-Whitney test. D: Example time course images of cell response to LMB treatment (120 min) and subsequent TNFα treatment. Scale bar = 20 µm. E: Example cell fluorescence tracking showing quantification of response to LMB and subsequent TNFα treatment. Nuclear intensity is indicated by unbroken lines, total cell intensity is indicated by dashed lines. F: qRT-PCR measurement of SK-N-AS target gene expression response to TNFα treatment following LMB preincubation for 1 – 24 hours. Target gene expression was normalised to *PPIA*. N = 3/treatment/time. Two-way ANOVA, Dunnett’s multiple comparison test. G: Chromatin immunoprecipitation using anti-p65 antibody against binding sites in the promoter regions of indicated target genes, with or without LMB pretreatment (120 min) or TNFα treatment (130 min). Samples for IgG and polII immunoprecipitations were treated with TNFα for 130 minutes. N = 2/treatment. H: Schematic representation of model of IκBα action on target gene expression.

Excess nuclear IκBα could potentially act as a competitor molecule to p65 action on target gene activation after TNFα treatment by preventing p65 binding to target gene promoters. We therefore investigated the effect of deliberately increasing IκBα nuclear concentration by LMB treatment prior to TNFα treatment. IκBα clone B cells transduced with p65-mCherry were treated for two hours with LMB and imaged by time lapse confocal microscopy (Figure 4D). Both the labelled IκBα and p65 proteins were observed to accumulate in the nucleus (Figure 4D and 4E; solid green line). Upon stimulation with TNFα, the remaining cytoplasmic IκBα was rapidly degraded, resulting in release of p65 to translocate to the nucleus. Notably, IκBα-eGFP which had already translocated to the nucleus in the presence of LMB was not degraded by stimulation with TNFα. This confirms that nuclear IκBα is protected from cytoplasmic signalling (Rodriguez et al., 1999). A transcriptional cycle was activated, evidenced by rise in total IκBα-eGFP levels, and newly synthesised IκBα-eGFP rapidly localises to the nucleus, but does not export p65 to the cytoplasm due to the continued presence of LMB (Figure 4E). Despite the continued presence of p65 in the nucleus no further IκBα-eGFP is detectably produced, indicating that p65 is now transcriptionally inactive. This may be due to IκBα complexing with p65 and removing it from the IκBα BAC transgene (and other target genes), although we cannot discount a role for dephosphorylation and inactivation of p65 through post-translational modification.

We hypothesised that increasing the concentration of IκBα could result in dampened downstream p65 target gene activation due to either (i) competitive binding by elevated nuclear IκBα to nuclear translocated p65, preventing binding and activation of target genes, or (ii) elevated cytoplasmic IκBα resulting in incomplete IκBα degradation in response to TNFα treatment, reducing the level of free p65 available for nuclear translocation. To test whether elevating nuclear IκBα levels can repress p65 target gene activation, we treated WT SK-N-AS cells with LMB to perturb the nuclear:cytoplasmic IκBα equilibrium and result in accumulation of IκBα and p65 in the nucleus. Following a range of periods of LMB treatment, we stimulated cells with TNFα for 130 minutes and examined the impact on downstream gene expression. After LMB pre-incubation and TNFα stimulation, mRNA was extracted and target gene expression was analysed by qRT-PCR. LMB treatment for up to 24 hours did not result in target gene activation despite the nuclear accumulation of p65 (Figure 4F), which corroborates previous findings (Sung et al., 2009). TNFα treatment in the absence of LMB resulted in strong target gene activation; this was significantly abrogated in cells subjected to pre-incubation with LMB, resulting in a marked reduction in target gene expression for all analysed targets after as little as one hour of LMB pre-treatment.

It is known that p65 activates target genes by binding NF-κB response elements (REs) in promoters, which facilitates the recruitment of transcriptional machinery (Zhang et al., 2017). If nuclear IκBα competitively prevents p65 binding to target sites in the genome, we would expect to see reduced p65 occupancy on NF-κB REs. Previous work in mouse B cells has found that increased nuclear occupancy of IκBα, in this study a result of nuclear export sequence (NES) mutation, reduced NF-κB recruitment to κB-containing probes on electrophoretic mobility shift assay (Wuerzberger-Davis et al., 2011). We used Chromatin Immunoprecipitation (ChIP) to investigate recruitment of p65 to well-characterised NF-κB REs in target gene promoters in WT cells. TNFα treatment for 130 minutes resulted in strong recruitment of p65 to all target gene promoters in the absence of LMB (Figure 4G). Interestingly, we also detected a low level of recruitment of p65 when cells were treated with LMB alone, indicating that nuclear p65 retains some capacity to bind target promoters whilst transcriptionally inactive. Following pre-treatment with LMB, 130 minutes of TNFα treatment did not result in p65 binding to target promoters (Figure 4G). This indicates that disrupting the equilibrium between nuclear and cytoplasmic localisation has a potent effect on gene activation. Overall, this supports our hypothesis that increasing the nuclear concentration of IκBα results in dampened downstream p65 target gene activation due to competitive binding by nuclear IκBα to translocated p65, reducing target gene activation (Figure 4H).

### An *in vivo* BAC IκBα overexpression model confirms the potency of nuclear IκBα to repress inflammatory gene activation

We used the IκBα-eGFP BAC to generate a transgenic mouse line to test this *in vivo*. Our transgenic line contains the IκBα-eGFP BACs at a single integration site on chromosome 8, with an estimated integration copy number of 2-6 (Figure S4). In primary fibroblasts from these mice, smFISH analysis indicates low IκBα transcript levels in the absence of treatment, with an induction after treatment with TNFα (Figure 5A). Transcript expression level appears lower than observed in our BAC clonal cell lines, and more comparable to WT SK-N-AS cells, likely as a result of the low copy number integration.

**Figure 5.**
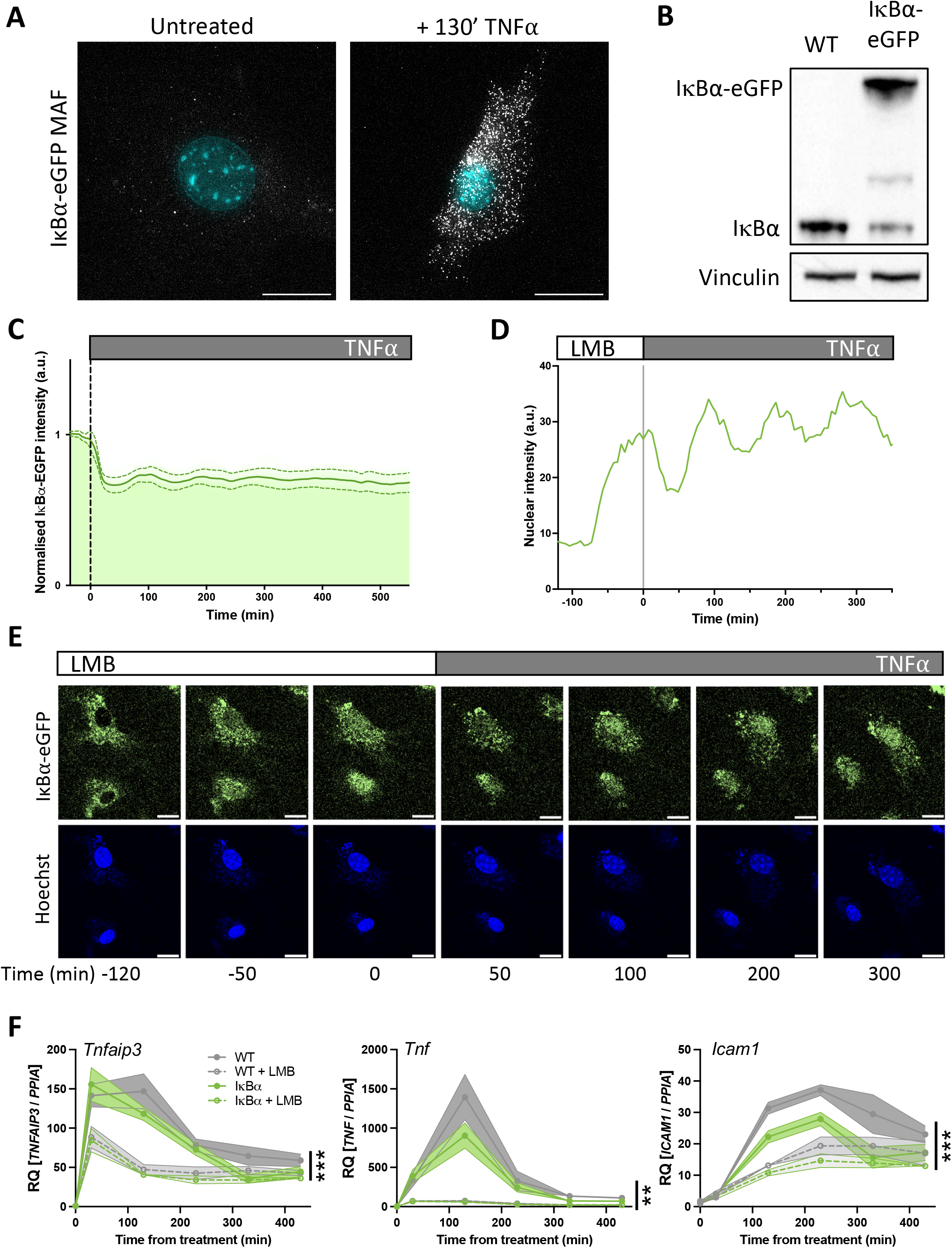
Primary mouse cell target gene expression is sensitive to nuclear IκBα accumulation. A: smRNA-FISH quantification of RNA transcripts in mouse adult fibroblast (MAF) cells. B: Western blot quantification of IκBα protein in MAF cells. C: Quantification of live cell nuclear fluorescence in IκBα-eGFP MAF cells in response to TNFα treatment. N = 16 cells. D: Example cell fluorescence tracking of live cell nuclear fluorescence in IκBα-eGFP MAF cells in response to LMB treatment. E: Example time course images of MAF response to LMB treatment (120 min) and subsequent TNFα treatment. Scale bar = 20 µm. F: qRT-PCR measurement of MAF gene expression response to TNFα treatment following LMB pretreatment for 120 min. Target gene expression was normalised to *Ppia*. N = 3/treatment/time/genotype. Three-way ANOVA with Geisser-Greenhouse correction; effect of LMB treatment is indicated to right of curves.

Western blot analysis showed a detectable band at native IκBα size in both WT mice and IκBα-eGFP BAC mice. Expression level of untagged endogenous IκBα appeared reduced compared to WT mice (around 30% of WT level, contrasting with the almost complete loss seen in the clonal cell lines), and we detect a strong higher molecular weight IκBα-eGFP band in the transgenic mice, resulting in total IκBα levels being around 1.8 times higher overall (Figure 5B). Upon TNFα treatment of the IκBα-eGFP BAC fibroblasts we detect oscillation cycles of degradation and resynthesis (with an initial peak at around 100 minutes after treatment; Figure 5C). We tested the effect of LMB treatment upon these cells and found, as with the clonal cell lines, that LMB treatment resulted in nuclear accumulation of the IκBα-eGFP fusion protein within 2 hours (Figure 5D-E). We see some residual oscillation in IκBα-eGFP nuclear intensity, which may indicate that inhibition of p65 activity as a result of nuclear IκBα accumulation is not complete in these cells, possibly due to lower levels of IκBα protein expression. This would result in reduced transcription of target genes, but not complete suppression.

To test this, we investigated the impact of LMB treatment and/or IκBα overexpression on the target genes *Tnfaip3, Tnf* and *Icam1* by RT-qPCR. IκBα overexpression resulted in reduced activation of *Tnf* and *Icam1*, but not *Tnfaip3*, after TNFα treatment. When cells were pre-treated with LMB, fibroblasts derived from both WT and IκBα-eGFP BAC mouse models exhibited lower activation of all three targets, with a pronounced loss of *Tnf* gene activation. These data corroborate the findings in BAC cell lines, showing that nuclear IκBα can have a suppressive effect upon NF-κB activation of target genes.

## Discussion

We have used a stably-integrated BAC reporter construct to investigate the dynamic response of the canonical NF-κB signalling proteins to cytokine stimulation in human and mouse cells. We find that, across a range of expression levels, p65 and IκBα reporter proteins show robust oscillatory dynamics. However, although we find protein dynamics in these systems to be insensitive to reporter overexpression, we find the transcriptional response of well-characterised NF-κB target genes to be broadly suppressed. This suppression is IκBα dose-dependent, and can be partially mitigated by overexpression of p65. We hypothesise that target gene transcriptional reduction results from increased levels of nuclear IκBα, and show that artificial increase of nuclear IκBα level, by treatment with the nuclear export inhibitor LMB, similarly suppresses target gene expression.

The NF-κB-IκBα signalling interaction is considered a canonical example of negative feedback regulation (Prescott et al., 2021). In our reporter model, we see high levels of IκBα transcript in basal and treated conditions (Figure 1B-D, Figure S1). As IκBα is a prominent negative regulator in the NF-κB signalling system, high transcript availability might be expected to shorten the interval between cytokine-induced NF-κB activation and subsequent repression; however, here we observed no effect on NF-κB dynamics. This indicates the control of oscillatory dynamics is decoupled from transcription, and the ‘inhibitory’ arm of the oscillation cycle (i.e. IκBα re-synthesis) is instead controlled at the post-translational level. Our previous work, using pulses of TNFα treatment, found the refractory period between activating cytokine treatment and induction of IκBα-mediated repression to be determined by a predicted enzymatic activity upstream of IKK activation (Adamson et al., 2016).

One can hypothesise that despite high levels of IκBα transcript, no new protein can be synthesised until deactivation of IKK occurs, and thus control of IKK activity may modulate oscillatory period (Fagerlund et al., 2015). In line with this, we have previously shown that manipulation of the level of *TNFAIP3*/A20 alters oscillatory period (Harper et al., 2018) and the refractory period between pulses of stimulation (Adamson et al., 2016). Other computational and experimental analyses have also found that NF-κB-inducible expression is not sufficient for effective negative feedback, and identified nuclear import and export of IκBα as key determinants of the dynamics of post-induction repression (Fagerlund et al., 2015). Our measurements find nuclear import rate to be comparable for both reporter cell lines, irrespective of protein expression level (Figure 4B-C), meaning this parameter does not alter oscillatory period in our reporter cells. More explicit modelling of the effect of nuclear import/export rates on oscillatory dynamics found persistent NF-κB translocation even in the presence of high levels of IκBα (Korwek et al., 2016). Taken together, these studies suggest IκBα is not the master driver of period, but a downstream effector, and period is controlled through IKK signalling and other regulatory feedbacks.

IκBα-deficient mice typically die shortly after birth, displaying signs of skin inflammation and increased expression of inflammatory chemokines (Rebholz et al., 2007). This suggests one key role of IκBα is in preventing ‘excessive’ transcription of inflammatory target genes. We hypothesise that the clear suppressive effect of IκBα overexpression on NF-κB target gene transcription (Figure 3) can be attributed to the increased levels of nuclear IκBα present (Figure 4A). Accumulation of labelled and endogenous IκBα in the nucleus in the absence of cytokine treatment has previously been taken as evidence for a basal level of shuttling between the cytoplasm and nucleus (Carlotti et al., 2000, Johnson et al., 1999), meaning high overall IκBα level is likely to result in higher nuclear levels of IκBα. The elevated nuclear IκBα level acts as a potent competitor for translocating active p65, preventing or attenuating promoter binding and effective target gene activation. Biophysical studies have shown that formation of a ternary complex with IκBα decreases the affinity of NF-κB for DNA and promotes formation of the stable, high affinity NF-κB-IκBα complex, resulting in thermodynamically favourable ‘molecular stripping’ of NF-κB from target promoters (Alverdi et al., 2014) (Potoyan et al., 2016). Mutation of the PEST interaction residues of IκBα slows removal of NF-κB complexes from DNA and relocation of NF-κB from the nucleus (Dembinski et al., 2017).

Mechanisms which promote the nuclear localisation of IκBα have been found to have a repressive effect on NF-κB-regulated targets in a number of previous studies. Pre-treatment of cells with LMB protects IκBα from signal-responsive degradation (Rodriguez et al., 1999). This results in a dose-dependent reduction in binding of NF-κB to kB probe sites in electrophoretic mobility shift assays (EMSAs), and reduces activity in transcriptional reporter assays (Huang et al., 2000). We see a similar effect upon the transcription of endogenous target genes in this study (Figure 4F, 5F), also previously seen elsewhere (Sung et al., 2009). We used chromatin immunoprecipitation to confirm that LMB pre-treatment reduces p65 binding at endogenous target promoters (Figure 4G). Promoting IκBα nuclear localisation by other strategies has been found to have a similar repressive effect upon kB reporter transcription. Overexpression of the receptor kinase GRK5 promotes nuclear accumulation of IκBα as a result of physical interaction and translocation, causing reduced kB-luciferase activity and NF-κB binding to kB probes in response to TNFα treatment (Sorriento et al., 2008). Mutation of the IκBα N-terminal NES also results in nuclear accumulation of IκBα (Wuerzberger-Davis et al., 2011), and these NES mutant mice show reduced expression of some NF-κB subunits and other target genes in response to LPS stimulation, as well as defective B cell maturation and lymph node development, and alterations in T cell development.

We cannot rule out that IκBα may have other effects in this overexpression system, as non-NF-κB-dependent functions have previously been identified. A SUMOylated form of IκBα has been identified in the nucleus of keratinocytes, where it physically interacts with H2A, H4 and PRC2 components including SUZ12 (Mulero et al., 2013). In this context IκBα has a repressive effect on PRC2 target genes, including members of the *HOX* family; IκBα binding at these loci is released by TNFα treatment, resulting in swift upregulation. SUMOylated IκBα has also been detected in the intestinal crypt cells of adult mice, with binding detected at promoters by ChIP-seq (Marruecos et al., 2020). IκBα again associated with PRC2 components in these cells, and knock-out of IκBα resulted in impaired maturation to adult cell identity, improving regenerative properties in response to inflammatory challenge. The effects we observe on the canonical NF-κB target genes of Cluster 1 (Figure 3) is responsive to ‘rebalancing’ via increasing p65 levels; however, effects on other genes might reflect pleiotropic regulation by IκBα. Another limitation of our work is that we have not characterised the effect of perturbation of other inhibitors and feedback genes (*NFKBIB, NFKBIE, TNFAIP3* etc.) on NF-κB-dependent transcription. *NFKBIE* has been shown to influence signalling response heterogeneity, whilst *TNFAIP3* can modulate NF-κB response to physiological conditions (Harper et al., 2018, Paszek et al., 2010). *NFKBIB* overexpression in carcinoma cell lines results in reduced NF-κB binding to DNA and downregulation of target gene transcription, with tumour-suppressive effects (Phoon et al., 2016). The IκB orthologues have non-redundant functions (Clark et al., 2011); further investigation may help to segregate orthologue functionality, but is beyond the scope of this paper. IκBα regulation also occurs post-translationally, and therapeutic intervention with proteasome inhibitors has been considered for diseases involving prominent NF-κB dysregulation (Wang et al., 2020, Vrabel et al., 2019). Introduction of a super-repressor form of IκBα that is resistant to proteolytic degradation improves survival in septic shock mouse models (Choi et al., 2020), suggesting increased IκBα level can have potentially beneficial anti-inflammatory effects in some contexts.

Overexpression of a gene is a classic experimental approach to inform gene and protein function. There are a number of ways in which this can be achieved, including small exogenously delivered constructs (plasmids, lentivirus), or, more recently, via activation of the endogenous gene locus by CRISPR activation (Adli, 2018). However, these methods fail to recapitulate the endogenous regulatory context of the overexpressed gene, so may not accurately reproduce native dynamic properties and responses to extrinsic stimulation. Overexpressing target genes using BACs provides copy number perturbation and increased levels of protein, but maintains regulation of the transgene in a pseudo-genomic context. In this study we used cell line tools and an *in vivo* mouse model that overexpress a labelled IκBα gene from a BAC that responds normally to an inflammatory stimulus. We used these models to demonstrate that IκBα overexpression has a pronounced effect on inflammatory gene activation, mediated through elevated nuclear IκBα that competes for translocating p65. This results in a broad anti-inflammatory effect without compromising the dynamic NF-κB signalling response to an activating inflammatory cytokine.

## Acknowledgements

This work was supported by BBSRC LoLa grant BB/K003097/1 (PD, DGS, DAJ, PP, MRHW), BBSRC David Phillips Fellowship BB/101796/1 (JSB, PP) and European Union SysMedIBD FP7 Programme agreement 305564 (JSB, HE, DAJ, MRHW, ADA).

We thank and acknowledge the University of Manchester Biological Services Facility staff for animal care, and the Bioimaging, Genome Editing and Genomic Technologies Facilities for experimental support. We thank E Boyd for providing primary tissue samples. The Bioimaging Facility SMC microscopes used in this study were purchased with grants from the BBSRC, Wellcome Trust and University of Manchester Strategic Fund.

## Notes

### Competing Interest Statement

The authors have declared no competing interest.

