## Supplementary Figures for "Overexpression of IκBα modulates NF-κB activation of inflammatory target gene expression"

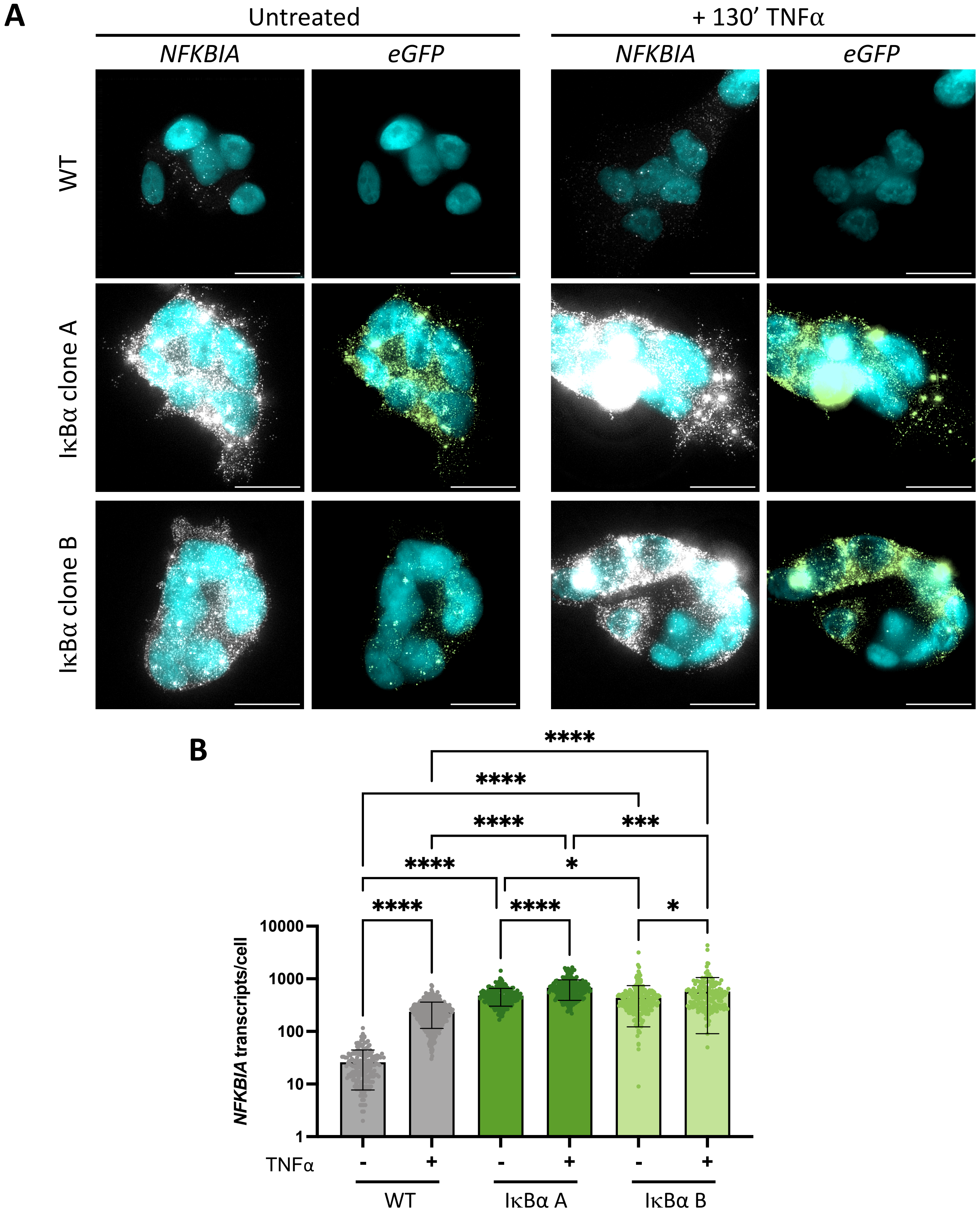


**Supplementary Figure 1. smRNA-FISH analysis of cell line response to TNFα treatment.**

A: smRNA-FISH detection of *NFKBIA* and *eGFP* transcripts with and without 130' TNFα treatment. Probe fluorescence is shown in white and green; DAPI counterstaining of nuclei is shown in cyan. Display settings are standardised for each fluorescence channel, resulting in signal saturation at bright spots. Scale bars = 20 μm.

B: Quantification of *NFKBIA* by smRNA-FISH detection in clonal cell lines with and without TNFα treatment. N = 200-400 cells/condition, imaged over 2-3 independent experiments. One-way ANOVA, Kruskal-Wallis test, Dunn's multiple comparison correction.

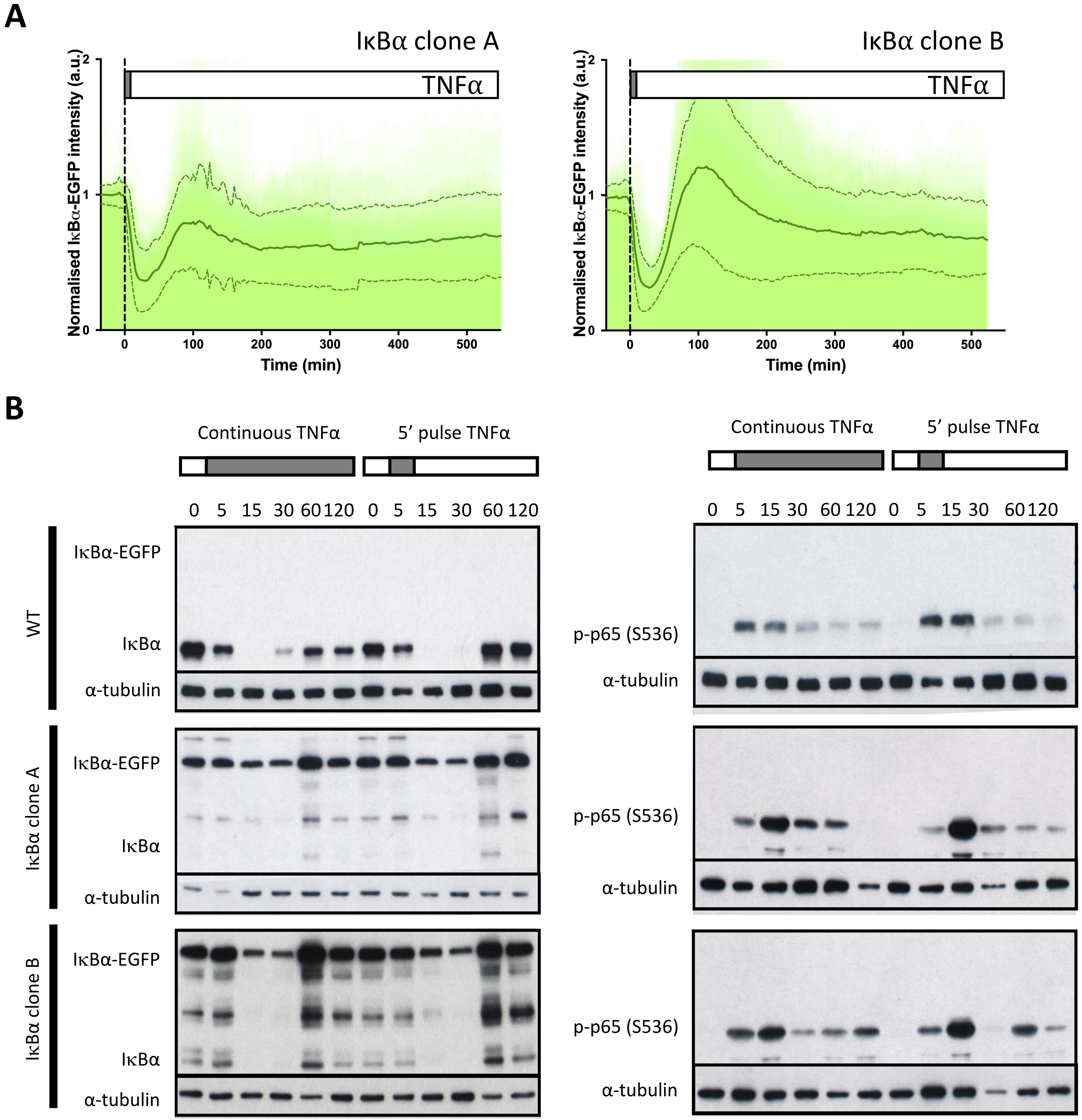


**Supplementary Figure 2. Dynamic response of IκBα-eGFP reporter cells to transient TNFa treatment.**

A: Clonal cell response to a 5 minute pulse of TNFα treatment. Individual traces are shown in pale green; population average +/- SD is shown in dark green. n = 140-150 cells imaged over at least 6 independent experiments.

B: Western blot analysis of protein response to TNFα treatment.

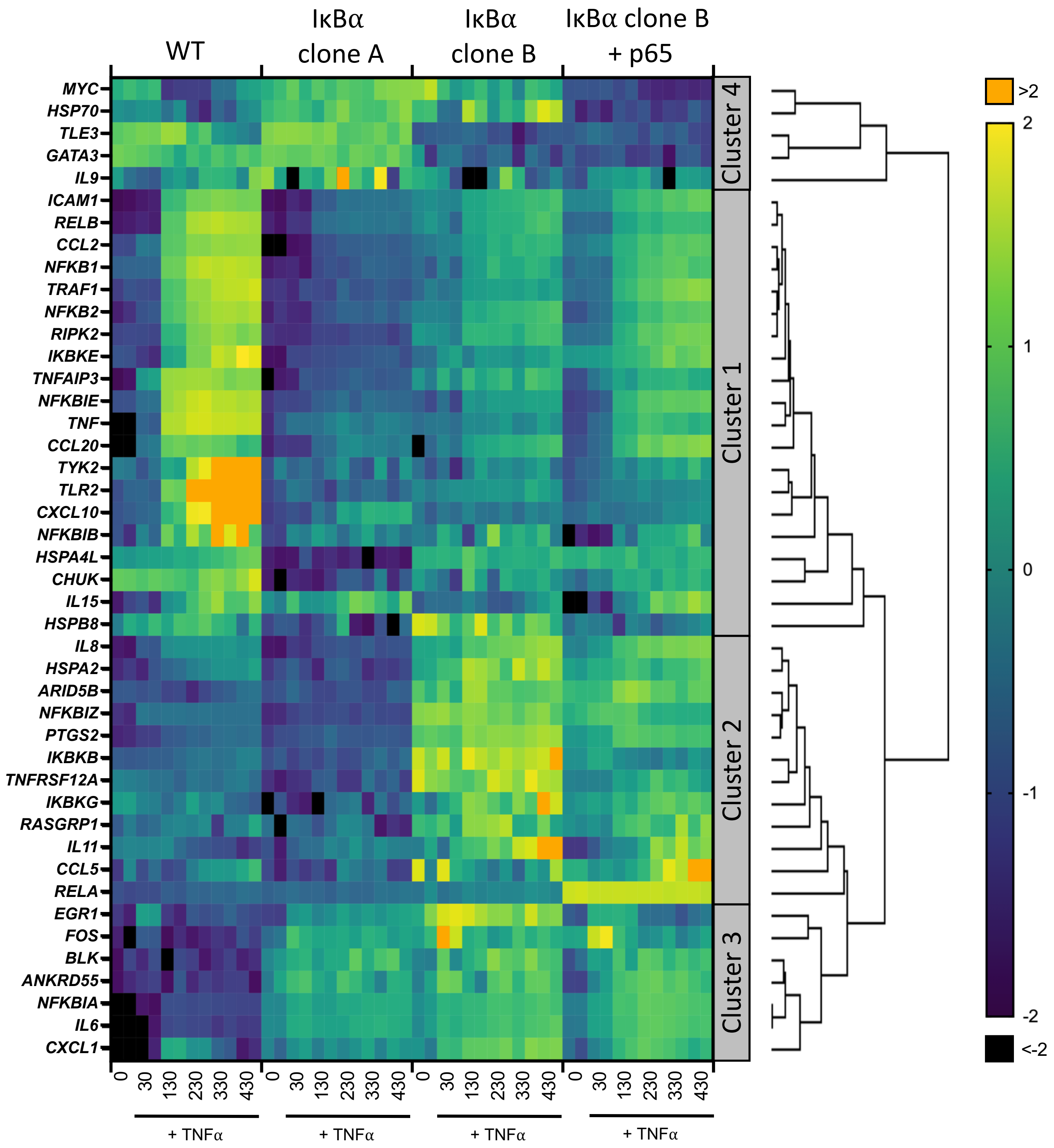


**Supplementary Figure 3. Nanostring analysis of cell line gene expression.** Heatmap representation of gene expression of clonal cell lines in response to TNFα treatment, as determined by Nanostring gene expression assay. N = 2/time/cell line. Expression is normalised to five housekeeping genes and internal control probes, then scaled to average gene expression level.

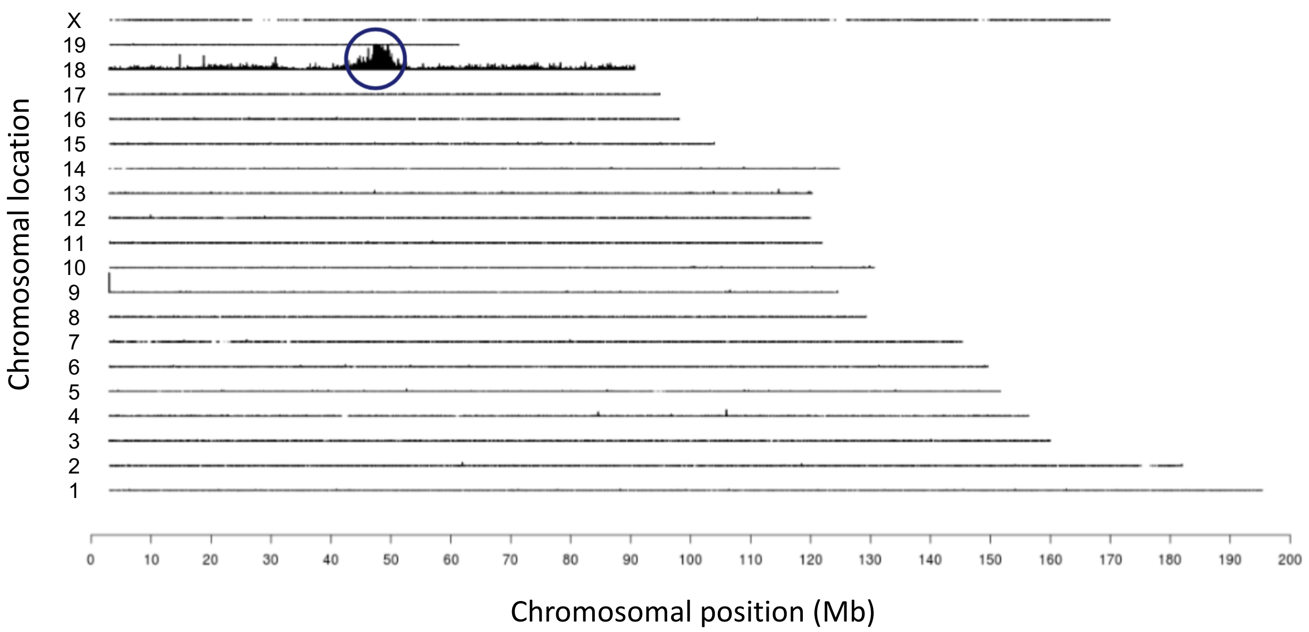


**Supplementary Figure 4. Copy number analysis of transgene integration.**
Targeted locus amplification and sequencing identified a single integration site of the IκBα-eGFP fusion BAC in mouse chromosome 18. Integration copy number was estimated as 2-6 copies based on read coverage.

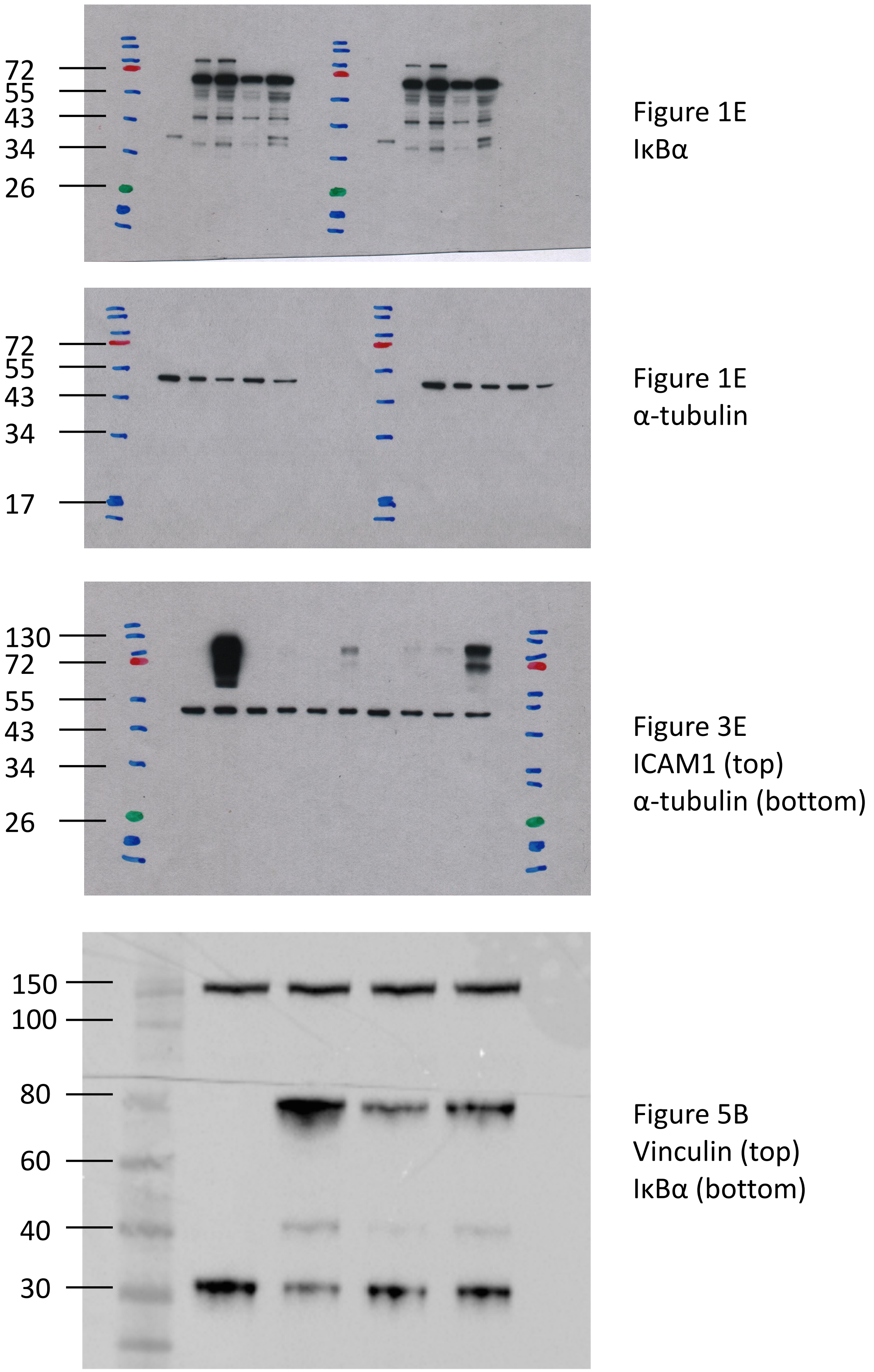


**Supplementary Figure 5. Unprocessed Western blot images.**
Protein ladder used: NEB P7712S (Figure 1E and 3E) or Thermo Scientific 84785 (Figure 5B).
